# Enzymes degraded under high light maintain proteostasis by transcriptional regulation in Arabidopsis

**DOI:** 10.1101/2021.10.03.462903

**Authors:** Lei Li, Owen Duncan, Diep R Ganguly, Chun Pong Lee, Peter A. Crisp, Akila Wijerathna-Yapa, Karzan Salih, Josua Trösch, Barry J Pogson, A. Harvey Millar

**Affiliations:** Frontiers Science Center for Cell Responses, Department of Plant Biology and Ecology, College of Life Sciences, Nankai University, 300071 Tianjin, China; ARC Centre of Excellence in Plant Energy Biology, School of Molecular Science, The University of Western Australia, 6009 Crawley, WA, Australia; Australian Research Council Centre of Excellence in Plant Energy Biology, Research School of Biology, Australian National University Canberra, Acton ACT 2601, Australia; CSIRO Synthetic Biology Future Science Platform, CSIRO, Action ACT, Australia; School of Agriculture and Food Sciences, The University of Queensland, Brisbane QLD 4072, Australia; Pharmaceutical Chemistry Department, Medical and Applied Science College, Charmo University, 46023 Chamchamal-Sulaimani, Kurdistan Region, Iraq

**Author notes:** **<u>Corresponding Authors</u>**, Lei Li -; A. Harvey Millar.

**Keywords:** Protein turnover, high light, protein homeostasis, transcription, translation

## Abstract

Photo-inhibitory high light stress in Arabidopsis leads to increases in markers of protein degradation and transcriptional upregulation of proteases and proteolytic machinery, but proteostasis is largely maintained. We find significant increases in the *in vivo* degradation rate for specific molecular chaperones, nitrate reductase, glyceraldehyde-3 phosphate dehydrogenase, and phosphoglycerate kinase and other plastid, mitochondrial, peroxisomal, and cytosolic enzymes involved in redox shuttles. Coupled analysis of protein degradation rates, mRNA levels, and protein abundance reveal that 57% of the nuclear-encoded enzymes with higher degradation rates also had high light-induced transcriptional responses to maintain proteostasis. In contrast, plastid-encoded proteins with enhanced degradation rates showed decreased transcript abundances and must maintain protein abundance by other processes. This analysis reveals a light-induced transcriptional program for nuclear-encoded genes, beyond the regulation of PSII D1 subunit and the function of PSII, to replace key protein degradation targets in plants and ensure proteostasis under high light stress.

## Introduction

Protein homeostasis (proteostasis) requires strictly controlled protein synthesis and degradation through coordinated gene expression, translational controls and protein degradation (Li et al., 2017; Millar et al., 2019). Protein turnover rates have been typically measured through a pulse-chase strategy by feeding plants or isolated organelles, radioactive precursors and monitoring the rates of appearance and disappearance of labelling (Sundby et al., 1993; Chotewutmontri and Barkan, 2020). The identification of PSII D1 subunit as the protein undergoing rapid turnover in chloroplasts, and the basis of photoinhibition under high light stress, was originally found using radioactive labelling of proteins in isolated chloroplasts (Ohad et al., 1984; Ohad et al., 1985; Sundby et al., 1993; Long et al., 1994). Using more recently developed discovery tools based on stable isotope labelling to measure turnover rates of many proteins, D1 was also noted as undergoing very rapid turnover in both Arabidopsis and barley, however these studies also identified other chloroplastic proteins being rapidly degraded under standard light conditions (Nelson et al., 2014b; Li et al., 2017). These studies have raised the prospect that a combination of direct or indirect photo-degradation targets may underlie photoinhibition and its consequences in plants (Li et al., 2018).

The degradation of soluble cytosolic proteins in plants typically occurs through the ubiquitin-proteasome pathway guided by selective ubiquitination of targeted proteins (Guo and Ecker, 2003). Plastids were commonly considered to be separated from this system by their membranes, thus relying on independent mechanisms of protein degradation (van Wijk and Kessler, 2017). However, recent research has uncovered an interconnection of these systems with specific plastid-localized proteins being tagged by ubiquitination to be degraded by the proteasome, a pathway termed Chloroplast-Associated Degradation (CHLORAD) (Ling et al., 2019). Chloroplasts damaged by UV exposure or over-accumulation of oxygen radicals are also degraded whole by globular vacuoles or by central vacuoles via selective autophagy (Woodson et al., 2015; Izumi et al., 2017). Specific protein degradation by selective autophagy has also been studied, but mainly for plastid stromal proteins such as RuBisCo (Michaeli et al., 2016). The proteolysis network inside chloroplasts works to differentially break down specific damaged proteins. CtpA and CtpA1 peptidase, CLP, DEG and FTSH family proteases have all been found or proposed, to play specialised roles in maturation, processing and cleavage of plastid-localized proteins (Gururani et al., 2015; van Wijk, 2015). As a consequence, there is ample opportunity for different rates of protein degradation to be initiated for specific plastid-localized proteins, which raises the question of how proteostasis is controlled when specific proteolytic processes are initiated.

High throughput studies have revealed rapid and robust changes in the metabolome, transcriptome, and proteome in plants during light and dark transitions or high light stresses (Vogel et al., 2014; Liang et al., 2016; Crisp et al., 2017; Huang et al., 2019; Schuster et al., 2020). Such studies typically confirm the lack of positive correlations between changes in steady state mRNA and protein abundance, *i.e*. compared with the rapid and robust changes in mRNA, protein abundances are often very stable and statistically significant changes in abundance are rare.

Here, we use high light induced photoinhibition to trigger protein degradation and explore the relationship between protein degradation rate, transcriptional responses, and protein abundance for enzymes that participate in the metabolic response to high light. In so doing, we have found new direct or indirect targets of photodamage in plants and shed light on how transcriptional processes counteract protein degradation to mask light-response changes in the proteome and enable proteostasis.

## Results

### High light leads to PSII photodamage and metabolic changes indicative of protein degradation

To analyse protein homeostasis under light stress, we performed a high light treatment of Arabidopsis plants aimed at inducing photoinhibition in conditions we could subsequently use to rapidly label proteins for analysis. We used a modified whole-plant growth chamber system (Kolling et al., 2015) and replaced the plexiglass lid with glass to increase light transmittance from an external LED light source to an Arabidopsis rosette inside (**Fig 3A**). Light intensity was held at 100 μE (standard light) or escalated to 500 μE (high light), and fluorescence pulse-amplitude-modulation (PAM) was utilized to evaluate PSII associated photochemical parameters inside leaves (**Fig 1**). After an hour of high light exposure, PSII parameters including Y(II) and Y(NPQ) showed significant changes under high light compared to standard light conditions (**Fig 1 A,B**) while Y(NO) remained steady under both light conditions (**Fig 1C**). This indicated 500 μE exceeded the maximum capacity of PSII, thus requiring energy dissipation through non-photochemical quenching. Dark adaptation could rescue the maximum quantum yield of PSII (Fv/Fm) after an hour of high light exposure but failed to restore Fv/Fm after 2, 5 or 8 hours of high light exposure (**Fig 1D**). Heat can contribute to non-photochemical quenching, so we measured the leaf surface temperature using an infrared thermometer. We could not detect statistically significant changes in leaf surface temperature on either the adaxial or abaxial leaf surface. The continuous room temperature air that was vented into the growth chamber during our measurements likely cooled the plant surface as shown in another recent high light stress study (Huang et al., 2019).

**Fig 1.**
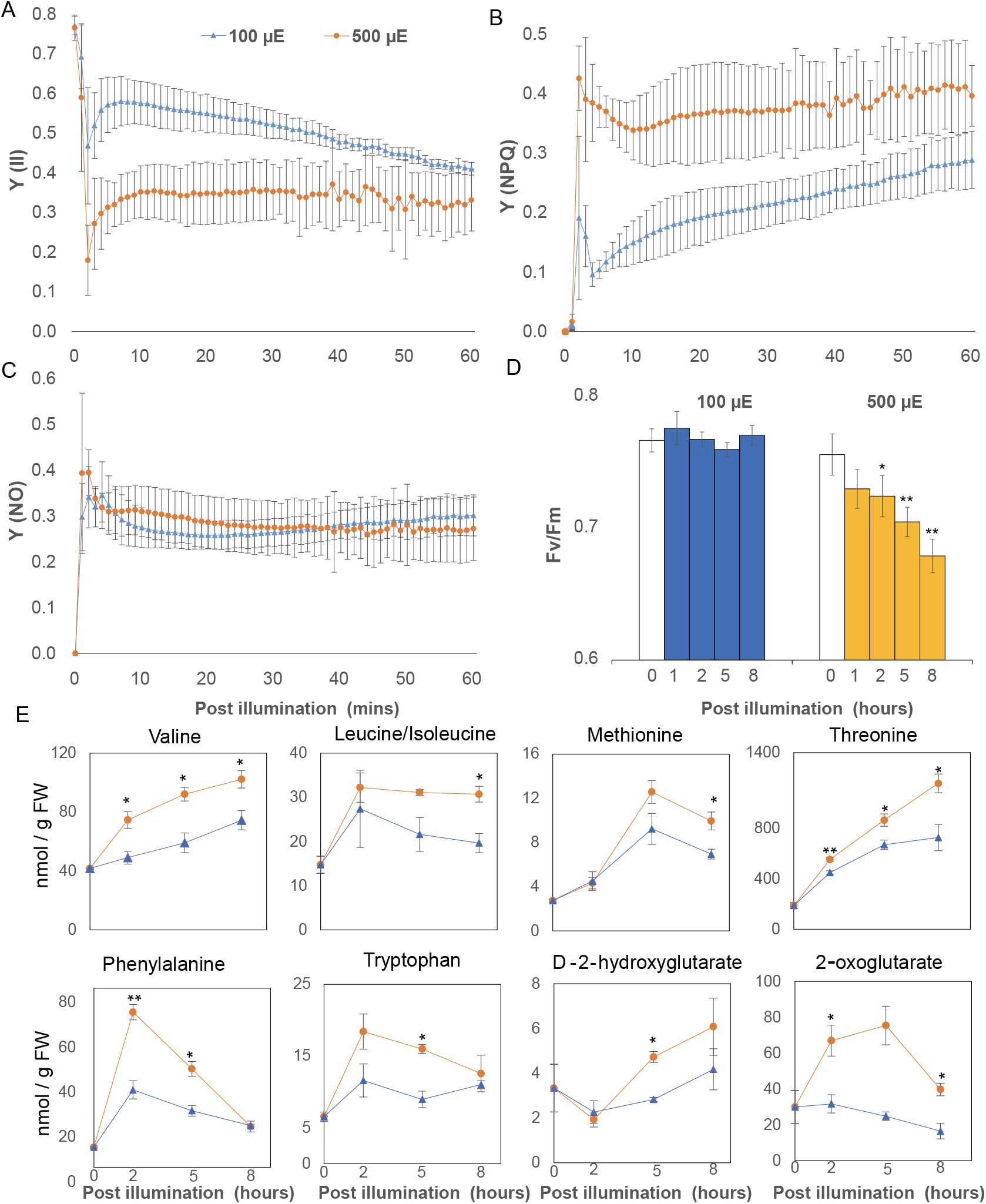
High light induced changes in photochemical responses and metabolite abundances in Arabidopsis. Arabidopsis in a growth chamber (**Fig 1 A**) were dark-adapted for at least 20 mins before being exposed to 100, and 500 μE LED light. Chlorophyll fluorescence measurements were transformed to three parameters that describe the fate of excitation energy in PS II, including Y(II)-quantum yields of photochemical energy conversion in PSII (**A**), Y(NPQ)-quantum yields of regulated non-photochemical energy loss in PSII (**B**) and Y(NO)-quantum yields of non-regulated non-photochemical energy loss in PSII (**C**). Arabidopsis plants were exposed to 100 and 500 μE LED light for 1,2,5 and 8 hours before their Fv/Fm values were determined by maxi-PAM (**D**). Specific amino acids (QQQ) and organic acids (Q-TOF) that increased in abundance in response to high light treatment (**E**). Error bars show standard deviations for photochemical parameters measurements (biological replicates n=4) and standard errors for metabolites measurements (biological replicates n=3). Statistical significance tests were performed with a student’s t test (**P<0.01, * P<0.05).

To measure the impact of photoinhibition of PSII on cellular metabolism, we measured amino acids, organic acids and sugar concentration in plants grown under standard and high light conditions (**Fig 1E, FigS1**). Six out of fourteen amino acids increased significantly (P<0.05) in abundance after high light treatment. Stress induced protein degradation products, including branched chain amino acids (Val, Leu and Ile) and aromatic amino acids (Phe, Trp and Tyr), were more abundant under high light. During plant stress, these protein degradation products serve as alternative respiratory substrates by being metabolized to D-2-hydroxyglutarate and 2-oxoglutarate (Araújo et al., 2011). Accordingly, both D-2-hydroxyglutarate and 2-oxoglutarate increased in abundance with high light treatment. These two metabolites are also known markers of high light dependent photorespiration (Kuhn et al., 2013). Aspartate abundance decreased significantly (P<0.05) after high light treatment (**FigS1**). However, threonine and methionine, which are biosynthesized from aspartate, and contribute to isoleucine biosynthesis (Hildebrandt et al., 2015), showed higher abundance after high light treatment. Sugars (sucrose, glucose, and fructose) and TCA cycle metabolites, other than 2-oxoglutarate, had comparable abundances over time between standard and high light conditions, with only citrate showing decreased abundance under high light.

### High light responses in the transcriptome correlate poorly with proteomic changes

To investigate the wider cellular response, we assessed changes in the transcriptome and the proteome under high light. Arabidopsis plants were transferred into the afore mentioned growth chamber and left overnight to acclimate before plants were treated for 2, 5 and 8 hours in standard or high light conditions, then harvested for protein or RNA. Total RNA sequencing (RNA-seq) detected 18,575 transcripts (**DataS1**) and quantitative proteomic experiments measured 1,548 protein abundances (**DataS2**).

Two hours of high light treatment led to the up or down-regulation of several hundred genes in Arabidopsis shoots **(Fig 2A)**. GO overrepresentation tests reveal enrichment of ontologies related to stress and unfolded protein responses (**DataS1**). By 5 h and 8 h of high light, there were several thousand differentially expressed genes **(Fig 2A)**. GO overrepresentation tests showed upregulation of genes encoding protein involved in RNA metabolism, translational, and nucleotide synthesis. At 8 h, up-regulated enrichment was also evident for proteolysis, proteasome and cellular catabolic processes (**DataS1**). While high light appeared to directly affect chloroplast fluorescence (**Fig 1**), transcriptional effects were mainly found in nuclear-encoded genes, with little evidence of changes in expression level of chloroplast encoded-genes (**Fig 2B**). Given the increase in amino acid abundances indicative of protein degradation (**Fig 1E**), and the GO enrichment analysis **(DataS1)**, we further investigated the expression of nuclear-genes encoding proteases and proteolytic machinery. We identified 30 nuclear-encoded genes in this ontological group that were differentially expressed in response to high light, most only reached significance after 8 h of treatment (**Fig 2C**). Of these, all the genes encoding for chloroplast-localised proteases were upregulated.

**Fig 2.**
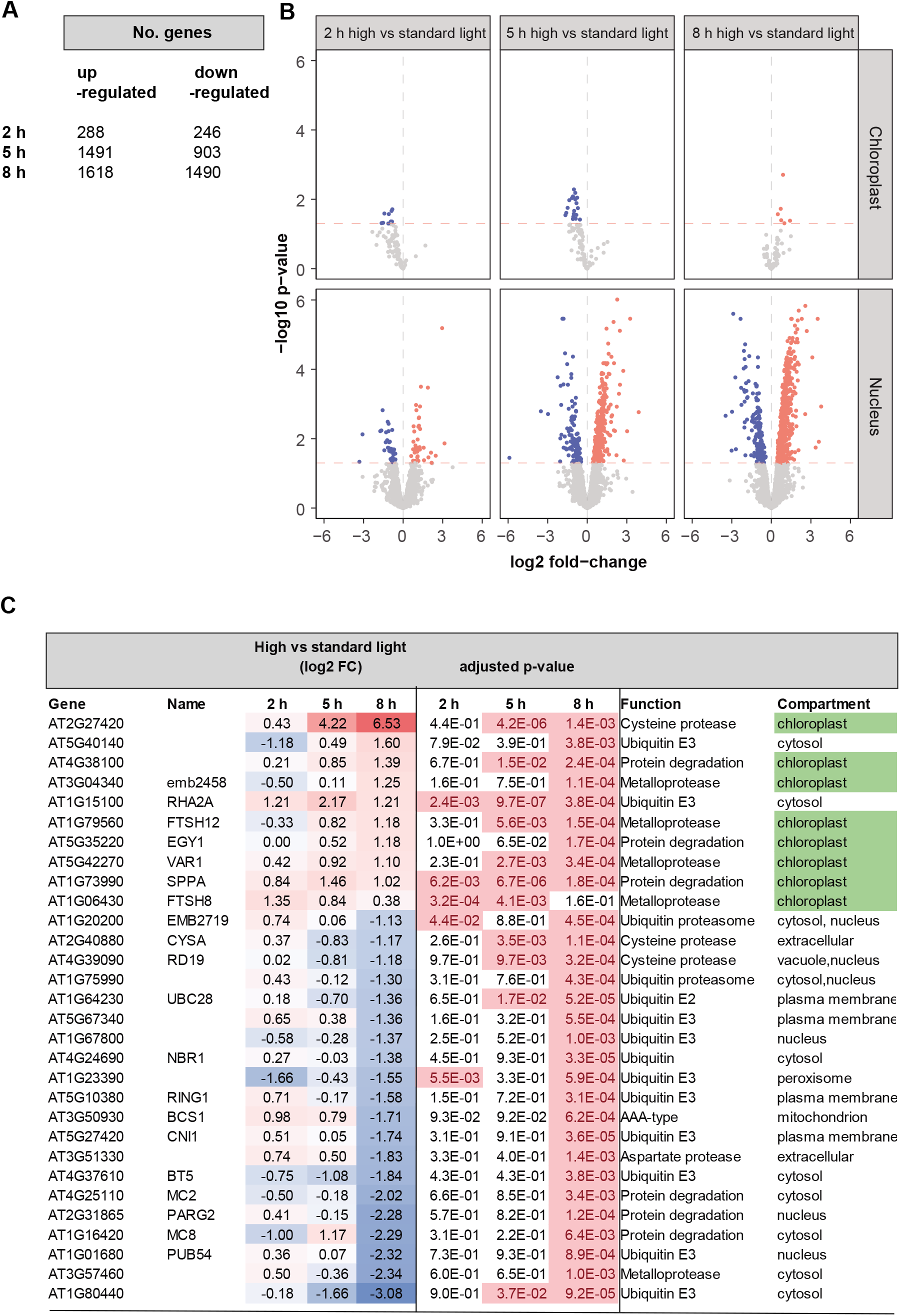
High light induced changes to the transcriptome in Arabidopsis shoots. Gene expression differences were induced by high light treatment of Arabidopsis shoot tissue. **(A)** Numbers of differentially expressed genes (adjusted p-value < 0.05) at 2 h, 5 h and 8 h of high light treatment. **(B)** Volcano plots of differential gene expression (horizontal red line denotes adjusted p-value < 0.05) of nuclear vs chloroplast-encoded genes at 2 h, 5 h and 8 h of high light treatment. Red and blue dots denote up- and down-regulated genes, respectively **(C)** Differential expression of genes encoding proteases and components of the proteolytic machinery, notably proteases in plastids (colored green).

To compare transcriptomic and proteomic responses, the abundance of proteins, measured across all samples, and their corresponding transcripts were extracted for PCA (**FigS2A,B**). While samples showed clustering by both time-point and light treatment based on transcript abundance, there was far less separation based on protein abundances. We also performed correlation analysis between the fold changes in protein and transcript abundance between high and standard light conditions (**FigS2 C-E**), which were found to be negligible (Pearson’s r: T2 = 0.08, T5 = 0.04, and T8 = 0.03).

### Direct measurement of protein turnover rates by partial ^13^CO2 labelling in Arabidopsis

To determine if proteins were being degraded in response to high light but then replaced, we sought to isotopically label new proteins and thus allow degradation of pre-existing proteins to be tracked by mass spectrometry. While we have previously used ^15^N labelling to assess protein degradation rates, the 4-6 h lag in this technique due to uptake by roots and translocation to leaves (Nelson et al., 2014b; Li et al., 2017) limited its utility to assess the impact of high light within 8 hours. ^13^CO_2_ fixation via photosynthesis is reported to allow rapid stable isotope incorporation in leaf amino acids and proteins (Ishihara et al., 2015; Ishihara et al., 2017). However, as the number of C atoms greatly exceeds N atoms in a tryptic peptide, the large mass shifts from ^13^C labelling greatly increase the complexity of the resulting peptide mass spectra (Nelson et al., 2014a). To minimize this effect, we calculated that lowering the ^13^C incorporation rate into Arabidopsis plants by supplementing air with 50% ^13^CO_2_ at 400 ppm could allow peptide mass spectra to be more readily interpreted. To conduct ^13^CO_2_ experiments, Arabidopsis plants were transferred into the growth chamber and left overnight to acclimate before labelling began in the morning. Arabidopsis plants were labelled for 2, 5 and 8 hours in standard or high light conditions, then harvested, and protein samples isolated and digested to measure isotopic incorporation and protein turnover rates. A representative mass spectrum of a tryptic peptide (ANLGMEVMHER) from a 2 hours sample shows the clear separation of the pre-existing peptide population (a typical natural abundance ^13^C labelling pattern with 1, 2 or 3 ^13^C atoms in the peptide) and a newly synthesized peptide population derived from the newly fixed ^13^CO_2_ (^13^C labelled pattern containing a median of 16 ^13^C atoms in the peptide) (**Fig 3A**). The technique itself is robust against the influence of differences in enrichment level because it measures labelled proteins of different enrichments as a group (Nelson et al., 2014b; Li et al., 2017). However, to determine if such differences exist between the two light regimes, we determined the ^13^C carbon content of the amino acids in the labelled peptide populations as described previously (Nelson et al., 2014b; Li et al., 2017). There were no differences in ^13^C enrichment between standard and high light conditions over the time course *i.e*. 2, 5 and 8 hours (**FigS3**, median enrichment: Std-light: 29%, 25% and 30%; H-light 29%, 27% and 34%). Mass spectra derived from peptides from three well-known rapidly turned over proteins (D1, THI1 and PIFI) showed that the ^13^C labelled peptide fraction (LPF) was one-third to one-half of the total peptide population under standard light conditions, indicating the rapid half-lives of these proteins (**Fig 3B**). A two-fold higher LPF was detected for D1 peptides under high light compared with standard light. Calculations of protein turnover rates over the time course measured ^13^C-derived protein turnover rates for 202 proteins in standard light and 269 proteins in high light (**DataS3**). The proteins measured had degradation rates (K_D_) of 0.15 to 10 per day representing a half-life range from 1.6 hours to 5 days.

**Fig 3.**
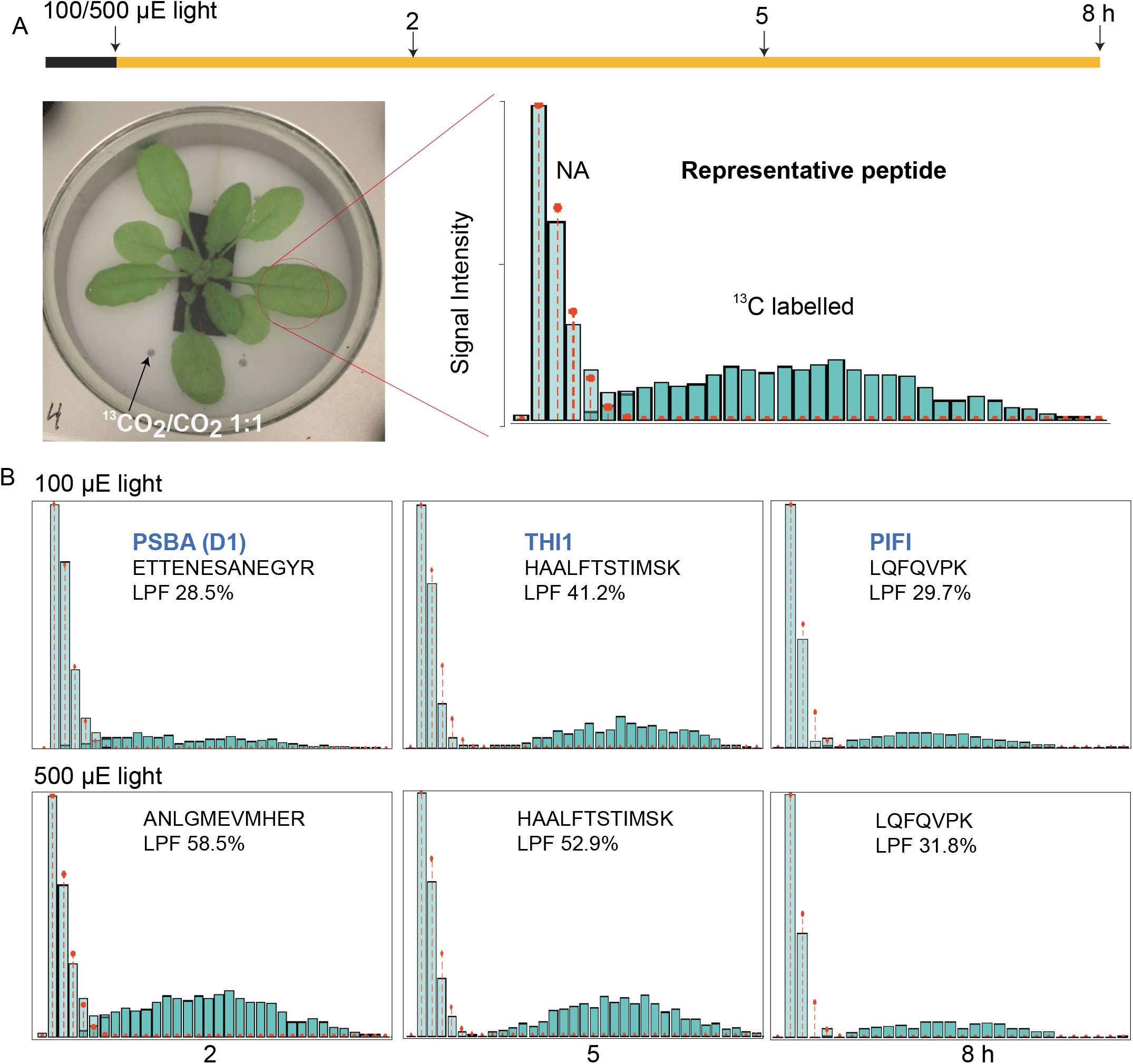
Measurement protein turnover rates in Arabidopsis shoots by partial ^13^CO_2_ labelling of the proteome. **(A)** Air containing ^13^CO_2_ was supplied at the end of night to a sealed growth chamber with a transparent glass lid allowing efficient light entry. Total proteins extracted from labelled shoots were analyzed by peptide mass spectrometry. A representative mass spectrum of one peptide from labelled shoot shows the natural abundance (NA) population and the new peptide synthesised using ^13^C labelled amino acids. (**B**) The mass spectra and calculated percentage labelled peptide fraction (LPF) for peptides derived from PSBA (D1;ATCG00020), THI1 (AT5G54770) and PIFI (AT3G15840) after 2, 5 and 8 hours of ^13^C labelling are shown. The natural abundance (NA) population is coloured light green and the newly synthesized peptide population is coloured dark green in each case.

### High light leads to faster turnover of photosynthetic proteins and associated enzymes in metabolic cascade reactions

Light stress can cause direct photodamage to D1 and change the turnover of proteins including D1 as well as other subunits of PSII and ATP synthase (Li et al., 2018; Chotewutmontri and Barkan, 2020). Using our ^13^C-derived protein turnover rates, we compared the difference in protein turnover rates between standard and high light conditions. This allows us to confirm expected, and discover new, direct targets of photodamage or proteins that are indirectly degraded under high light. Protein turnover rates of 140 proteins could be compared between two light conditions (**DataS4**). Compared to measurement under standard light conditions, 74 out of 140 proteins showed statistically significant changes in degradation rate under high light, 73 showed faster turnover, while one protein (protochlorophyllide oxidoreducase B-PORB, At4g27440) turned over more slowly. Placing the proteins with measured degradation rates in their functional, metabolic, and subcellular contexts shows the depth of impact that high light has on protein degradation rates in Arabidopsis rosettes (**Fig 4**). D1 (PSBA) showed the fastest rate of degradation overall and a 3-fold increase in degradation rate under high light, while other PSII subunits, PSB28 and PSBP, showed lower median degradation rates but still a 2-3 fold increase in degradation rate under high light. ATP synthase subunits (α, ε and b/b’) also showed significantly faster turnover under high light. A similar degree of degradation rate induction was seen for a series of molecular chaperones in the chloroplast and mitochondria, and also specific enzymes involved in the Calvin-Benson cycle (CBC) and glycolysis. All the major members of the malate dehydrogenase (MDH) family, which catalyze the malate shuttle between organelles, as well as thioredoxin and glutaredoxin linked enzymes in the chloroplast and mitochondria also degraded more rapidly under high light conditions.

**Fig 4.**
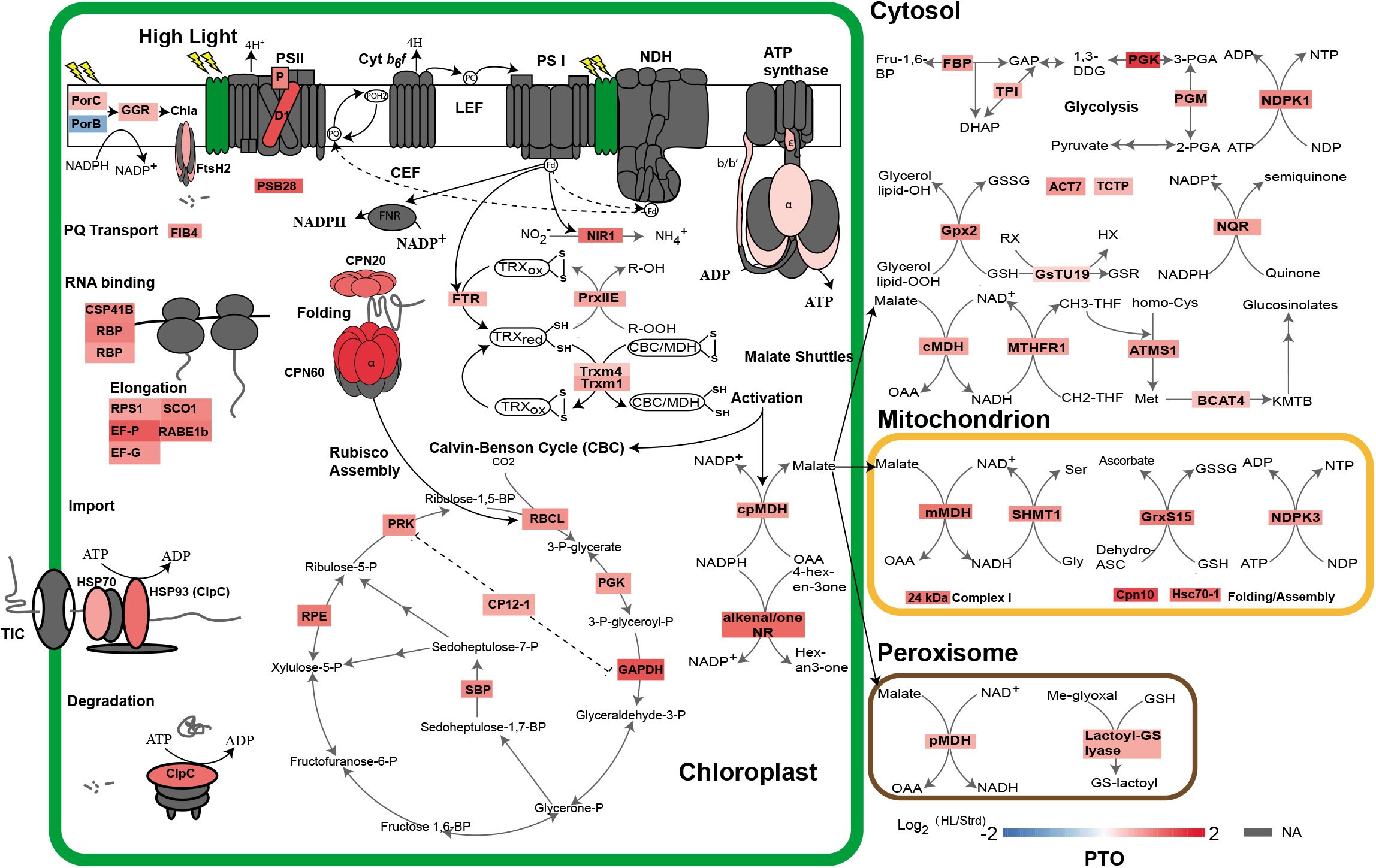
Changes in protein turnover rates in response to high light treatment. Changes in protein degradation rate were shown as log2 fold changes between high and standard light. For visualization, sixty-one proteins with annotated functions and localize in four major cellular compartments *i.e*. chloroplast, cytosol, mitochondrion and peroxisome were extracted from the set of 74 proteins with significant changes in rate. Protein subunits in photosynthetic complexes, import apparatus, chaperonin and protease were coloured according to the values of log2 fold changes (**Data S4**). Protein subunits with non-significant changes or non-available data were coloured grey.

Notably, there was no change in degradation rate of LHC-II, PSI, NDH or Cyt *b_6_f* subunits (**Fig 4, DataS4**). However, NIR1 and FTR, which take electrons from PSI Fd to reduce oxidized thioredoxin and nitrite, clearly degrade faster in high light. It was reported that reduced thioredoxin from FTR can serve as a reductant for activation of MDH and CBC, which links PSI Fd with metabolic enzymes in the chloroplast (Dai et al., 2000; Collin et al., 2003; Marri et al., 2009; Michelet et al., 2013; Guinea Diaz et al., 2020; Yu et al., 2020). Here we show that Trxm1 and Trxm4 turned over faster, as did chloroplast MDH and CBC enzymes, in response to high light. Furthermore, faster turnover of malate shuttles linked metabolic enzymes were observed. There was a faster turnover of MTHFR1, ATMS1 and BCAT4 in the cytosol that catalyse the reductive conversion of 5,10-methylenetetrahydrofolate (CH2-THF) to 5-methyltetrahydrofolate (CH3-THF), which then serves as a methyl donor for methionine biosynthesis and the following chain elongation pathway. In mitochondria, SHMT1, which catalyses the production of serine from glycine, degraded faster under high light. A number of enzymes involved in the consumption of NADPH, NADH, ATP, and glutathione also show faster degradation rates (**Fig 4, DataS4**). Examples of this group include PORC, which catalyses the conversion of protochlorophyllide to chlorophyllide, and geranylgeranyl (GG) chlorophyll a reductase-GGR, which catalyse the formation of chlorophyll *a* in the thylakoid membrane; cytosolic NDPK1 and mitochondrial NDPK3 that catalyses the production of nucleotides by consuming ATP; cytosolic NADPH:Quinone Reductase-NQR that converts quinone to semiquinone; and Gpx2, GsTU19, GrxS15 and Lactoyl-GS lyase that reduce oxidative metabolites by consuming glutathione. Taken together, we can see a clear pattern of faster turnover of enzymes involved in metabolic reactions responding to high light.

Beyond metabolic enzymes, faster degradation was also observed for a number of elongation factors, chaperonins, and proteases in response to high light (**Fig 4, DataS4**). Proteins in this group are essential for protein synthesis, folding, assembly, and degradation to maintain proteostasis. This group includes RNA binding proteins (CSP41B and RBPs) and elongation factors (EF-P/G, RPS1, SCO1 and RABE1b) in the chloroplast; cpHSP70 and HSP93-V (ClpC) that are involved in chloroplast protein import (Nakai, 2018); and ClpC that forms a protein complex with CLP protease to unfold selected proteins for degradation (van Wijk, 2015; Nakai, 2018); chloroplast CPN20 and CPN60 that form a protein complex for the assembly of RuBisCo (Aigner et al., 2017; Vitlin Gruber and Feiz, 2018); mitochondrial chaperonin CPN10 and mtHsc70-1 involved in electron transport chain protein complex assembly (Wei et al., 2019); and FtsH2 that forms a protein complex with FtsH1/5/8 and is involved in D1 degradation (Zaltsman et al., 2005; Nishimura et al., 2016).

### Transcriptional responses counteract increased protein turnover to maintain proteostasis of many major cellular enzymes

To further examine the 74 proteins exhibiting light-induced changes in degradation rate, we performed fuzzy k-mean clustering based on their changes in degradation rate and transcript abundance (**Fig 5A, DataS5**). This approach grouped the 74 proteins into three clusters. Proteins in clusters 1 and 3 had the same change in degradation rate, however, cluster 1 genes were up-regulated by high light whereas those in cluster 2 were down-regulated. Cluster 3 contained proteins with greater changes in degradation rate whose encoding genes were firstly up-regulated and then down-regulated by high light.

**Fig 5.**
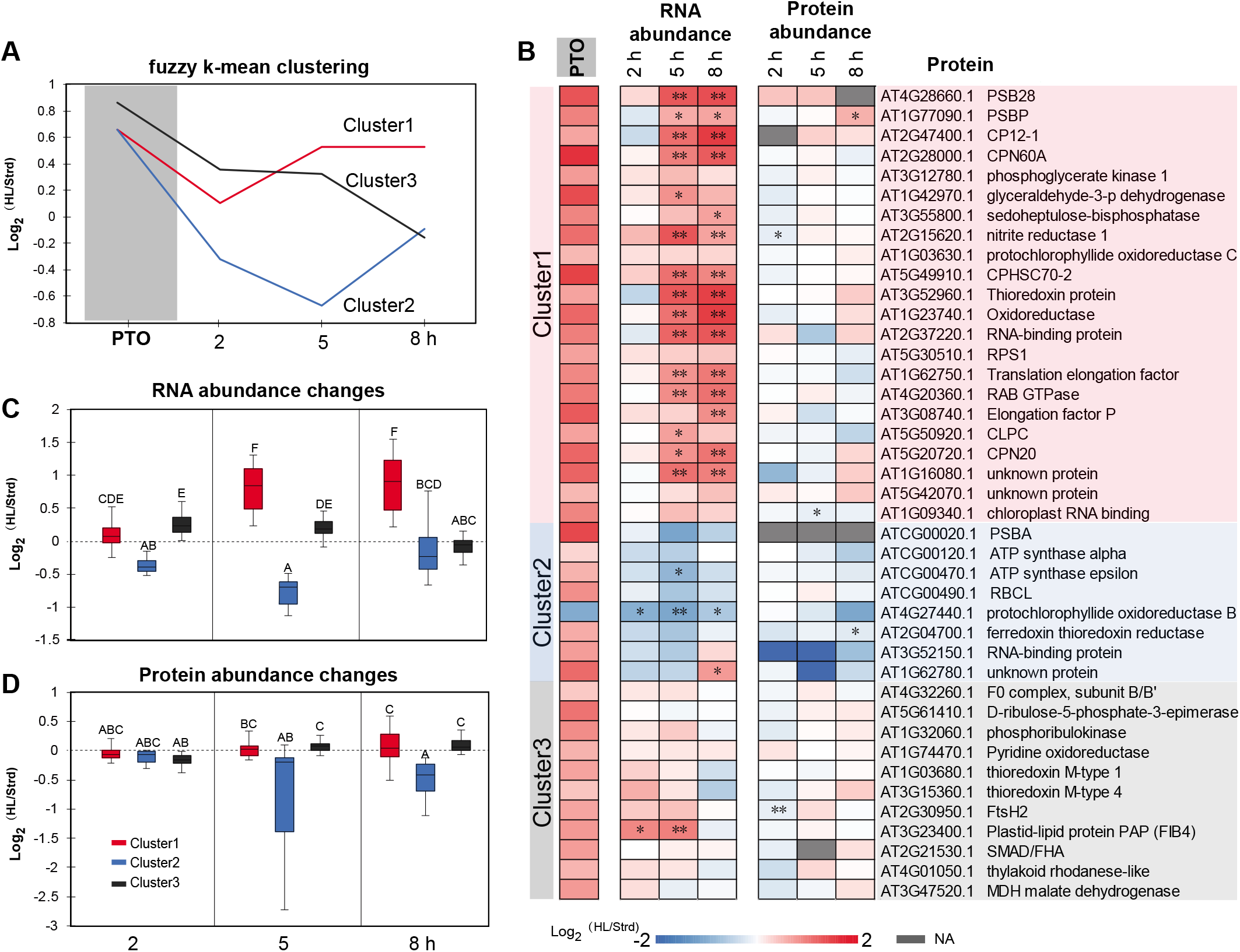
Changes in transcript and protein abundance for proteins with significant changes in protein turnover rate during high light treatment. Based on patterns of protein degradation and transcript changes, a fuzzy k-mean clustering method was utilized to cluster the 74 proteins with significant changes in protein turnover rate. Representative curves of the three clusters were plotted **(A)** and values of distance to centroid for specific proteins is provided in **Data S5**. Forty-one plastid proteins were extracted from the whole set to show their protein turnover rate alongside fold changes in transcript and protein abundance **(B)**. Boxplots of changes in transcript **(C)** and protein abundance **(D)** of each cluster over the time course are shown. PTO; change in protein turnover rate.

A total of 41 out of the 74 light-induced degradation set were chloroplast-localized proteins (**DataS5)**. To further investigate the coordination between protein turnover, and RNA and protein abundance in the chloroplast, we aligned the changes (log_2_ fold-change) in these traits between standard and high light conditions, then ranked them by functional categories in the three clusters (**Fig 5B**). In contrast to their faster turnover in response to high light treatment, chloroplast protein abundance for these proteins remained unchanged. We only found FtsH2 protease, RNA binding protein-At1g09340, and NIR1 that showed statistically significant abundance decreases after 2 or 5 hours of high light treatment (**Fig 5B**). PSII subunit PSBP even showed a statistically significant increase in abundance after 8 hours. This indicated proteostasis of fast turnover chloroplast proteins is maintained even after high light treatment for 8 hours. Transcriptional up-regulation for the genes encoding proteins in Cluster 1 and 3 masked their faster turnover, thus maintaining protein levels. Cluster 2 consisted of eight proteins, with half of them being encoded by the chloroplast genome. All chloroplast encoded proteins with measured turnover in this study, namely PSII D1, ATP synthase subunits (α and ε), and RuBisCo large subunit follow the Cluster 2 pattern (**Fig 5B**). Expression of these chloroplast-encoded genes are not induced by high light stress treatment, so their proteostasis in the face of increased protein degradation rates must be governed by post-transcriptional process.

To investigate the timing of coordination between transcription and proteostasis for chloroplast proteins, we plotted RNA and protein abundance changes by members in each cluster (**Fig 5C-D**). Consistently, smaller net changes were observed in protein abundance (statistical grouping A-C) than RNA abundance (statistical grouping A-E). Net protein abundances started to decrease after 2 hours of high light exposure. However, protein abundance for Cluster 1 and 3 started to recover while the abundance of members of Cluster 2 continued decreasing over the time course of high light treatment. It is evident that Cluster 1 and 3 complemented their faster protein turnover through enhanced transcription. A higher level of transcripts permitted continued protein translational to counteract light-induced protein degradation. In contrast, Cluster 2 were more inclined to drop in both transcription and protein abundance after high light treatment. Their proteostasis is likely to be recovered more slowly through post-transcriptional responses involving translational control.

## Discussion

### A multi-omics analysis reveals new targets of light-dependent protein degradation

It is well known that high light stress causes damage to the PSII reaction centre protein, D1, and leads to impaired PSII efficiency. In this study, we found high light tripled the degradation rate of D1 with a concomitant drop in PSII efficiency [Y(II)] (**Fig 1, DataS4**). We observed that although Y(II) dropped at the beginning of the high light treatment it gradually recovered to the level observed under standard light over the first hour of high light exposure (**Fig 1A**). This suggests Arabidopsis plants can cope with the increased turnover rate of D1 under high light by maintaining proteostasis and PSII function after a short time course of high light exposure. Consistent with this, a recent study utilizing ribosomal profiling and pulse labelling found that D1 photodamage can trigger recruitment of its mRNA to the ribosome to enhance D1 synthesis (Chotewutmontri and Barkan, 2020). This demonstrates that D1 degradation and synthesis are matched to maintain proteostasis for short-term high light acclimation. However, we found that Fv/Fm, an indicator of PSII maximum efficiency after dark adaption, declined after longer high light exposure. This suggests that longer periods of high light caused irreversible damage, from which Arabidopsis PSII efficiency cannot recover even after dark adaptation, likely due to the uncoupling of D1 degradation from its synthesis rate (**Fig 5**).

Beyond the D1 protein, we found high light significantly increased the degradation of another seventy-two proteins (**DataS4**). Protein degradation in our high light experiments is supported by our measures of the accumulation of amino acids through protein degradation (**Fig 1, FigS1**) and up-regulation of protease gene expression (**Fig 2**). To investigate how Arabidopsis coped with this enhanced degradation to maintain proteostasis, we investigated changes in transcript abundances between standard and high light conditions (**Fig 2,5**). Nuclear-encoded genes encoding proteins with high turnover rates (clusters 1 and 3) demonstrated transcriptional responses that masked protein turnover changes, resulting in proteostasis under high light (**Fig 5 A-C**). These strong correlations between faster protein turnover and higher transcript abundances help explain the purpose of high light triggered transcript induction without apparent protein abundance changes.

In contrast, chloroplast encoded genes (D1, Rubisco large subunit, and ATP synthase subunits) do not respond to high light at the transcriptional level, and their RNA levels even dropped to some extent. This limited transcription response in the chloroplast under light stress was also reported in tobacco (Schuster et al., 2020). It appears that chloroplast-encoded genes largely rely on post-transcriptional controls to counteract rapid protein turnover under high light. Previous studies focusing on *in vitro* or *in vivo* chloroplast translation observed translation elongation rate stimulated by light (Muhlbauer and Eichacker, 1998; Trebitsh and Danon, 2001; Chotewutmontri and Barkan, 2016; Chotewutmontri and Barkan, 2020). The activation of protein synthesis by elongation is also supported by faster turnover of different RNA binding proteins and elongation factors in this study (**Fig 4**). For short-term high light exposure, rapid protein synthesis from translation elongation can complement rapid protein degradation due to photodamage to maintain proteostasis. However, we found chloroplast translation failed to keep pace with protein degradation after a longer periods of high light exposure. This is supported by the failure of dark adaptation to recover PSII (**Fig 1D**) and the tendency towards protein abundance decreases after a longer high light exposure (**Fig 5D**). Recently, a salvaging strategy to circumvent inefficient chloroplast translation by expressing D1 protein from the nuclear genome was found to enhance Arabidopsis, tobacco and rice performance under stress conditions (Chen et al., 2020). It would be attractive to perform a wider salvaging operation involving other photodamage targets discovered in this study to maintain their proteostasis under high light or other stresses.

### Many fast turnover proteins are unaffected by high light

We found that high light does not affect turnover rates for nearly half of the 140 proteins that we could assess between standard and high light conditions (**DataS4**). Some of these are rapidly turnover proteins such as PIFI (Post-Illumination Chlorophyll Fluorescence Increase) and CCD4 (Carotenoid Cleavage Dioxygenase 4). PIFI is an ancillary subunit of the chloroplast NDH complex, and we previously proposed PIFI’s rapid turnover could relate to the putative role of the NDH complex in photoprotection (Li et al., 2017; Li et al., 2018). But our high light data suggests the control of PIFI turnover is independent of light stress. CCD4 is a plastoglobuli-localized enzyme that cleaves carotenoids, such as β-carotene (Gonzalez-Jorge et al., 2013; van Wijk and Kessler, 2017). Its degradation was proposed to associate with a plastoglobuli M48 peptidase PGM48 (Bhuiyan et al., 2016). *In silico* modelling of CCD4 suggests it has lower stability compared with other members of the CCD gene family (Priya et al., 2017). Its rapid turnover may reflect its suborganelle location, which is distinct to the other CCDs or this modelled intrinsic lower stability rather than light stress. Rapid degradation of other proteins, such as CML10, THI1, GRP2, BAM3, show only small rate changes in high light. It is probable, at least for these proteins, that their rapid turnover rates are due to their function, sequence, protein domains, or cellular location rather than light stress (Li et al., 2017).

We also observed that the rapidly turning over enzyme, protochlorophyllide oxidoreductase (PORB) exhibited a significant slowing of its degradation rate under high light (**Fig 4**). Protochlorophyllide oxidoreductase is a light-activated enzyme, which catalyzes the transformation of protochlorophyllide to chlorophyllide. In barley and rice, there are two isoforms of protochlorophyllide oxidoreductases whose expression are regulated differently by normal and high light (Holtorf et al., 1995; Lebedev and Timko, 1999; Sakuraba et al., 2013). In Arabidopsis, there are three protochlorophyllide oxidoreductases namely PORA, PORB and PORC (Gabruk and Mysliwa-Kurdziel, 2015). Repression of Arabidopsis PORB gene expression by light has been reported (Hoecker and Quail, 2001; Matsumoto et al., 2004). In this study, we also found *PORB* gene expression is repressed after high light treatment. In contrast, PORC showed faster protein turnover in high light and slightly induced gene expression. It is conceivable that PORC plays a specific role in chlorophyll biogenesis under high light conditions. For PORB and PORC, transcription plays a key role in maintaining proteostasis, and their protein turnover rates appear to be responsive to changes in transcript abundance.

### Metabolic explanation of increased protein turnover rates

The turnover of D1 is typically explained as a response to photo-inactivation of the protein. Research suggests that photodamage to PSII may involve the disintegration of the Mn^2+^ centre in PSII that leads to an energy imbalance and, a so far ill-defined oxidative damage of residues in D1. Loss of D1 impairs PSII function and leads to cleavage of the damaged D1 subunit by proteases in a two-step model (Kato et al., 2015). Turnover of another rapidly degrading protein, thiamin synthase (THI1), is explained by its suicide mechanism that means the enzyme has a single catalytic cycle before it is inactivated and needs to be replaced (Chatterjee et al., 2011; Joshi et al., 2020). Recently we showed across a wide range of enzymes in Arabidopsis, yeast and bacteria, that the number of catalytic cycles until replacement varied according to the chemical risk of the reaction they undertook, including enzymes with photoactivatable substrates or with reactive oxygen producing roles in metabolism (Hanson et al., 2021). It is evident from our protein turnover measurements that high light leads to faster degradation of PSII D1, PSB28, PSBP, PORC and GGR, which all catalyse light activated reactions (**Fig 4, DataS4**). PSI was also activated by high light, yet seemingly its rate of protein degradation was unaffected. FTR and NIR1, which take electrons from PSI Fd to reduce thioredoxin and nitrite, turned over faster. Thioredoxins can serve as reductants to activate CBC and MDH catalyst activities (Nikkanen and Rintamaki, 2019). Moreover, transient overproduction of NADPH and ATP as substrates may further accelerate the usage of CBC and MDH enzymes, and also accelerate their turnover. MDH activation in the chloroplast acts as a stimulus to malate circulation to the cytosol, mitochondrion, and peroxisome in so-called malate shuttles of reductant from sites of synthesis to cellular sinks (Selinski and Scheibe, 2019). In terms of risk, high light and photoinhibition is likely to lead to increased ROS production and an elevated need for shuttling of reductant out of the plastid to other cellular compartments. The increased turnover of redox shuttling systems, namely glutathione and thioredoxin linked systems and the MDH enzymes involved in malate shuttles throughout the cell **(Fig 4)**, may be due to increased flux through these pathways and thus a consequence of an increased rate of wear-out damage of these enzymes (Hanson et al., 2021; Tivendale et al., 2021).

### Conclusion

We have discovered a range of proteins with enhanced rates of degradation in response to high light. Light-activated electron transport pathways and metabolic fluxes likely stimulate the usage of metabolic enzymes and accelerate their degradation. Potential protein targets of photodamage, many of which are chloroplast-localized, have been revealed, and a differential role of nuclear and plastid transcriptional control to maintain proteostasis has been highlighted.

## Methods

### Arabidopsis plants preparation and ^13^CO_2_ labelling

*Arabidopsis thaliana* accession Columbia-0 plants were grown under 16/8-h light/dark conditions with cool white T8 tubular fluorescent lamps 4000K 3350 lm (Osram, Germany) with the intensity of 100–125 μmol m^−2^ s^−1^ at 22 °C. Arabidopsis plants were grown in soil pots for 21 days until they reached leaf production stage 1.10 (Boyes et al., 2001). Shoots of Arabidopsis at the leaf production stage 1.10 were positioned into the sealed growth chamber with the soil pots kept underneath (**Fig 3**). Six tandem growth chambers were supplied with air at a continuous flow rate 6 L/min and kept overnight before the labelling experiment (T_0_). A homemade water column was connected to the air hose to keep the air humidity inside the growth chamber. A commercial LED (Heliospectra) was used as the light source for the labelling experiment, and the light spectra was set as (420nm-250, 450nm-638, 530nm-750, 630nm-1000, 660nm-250 and 735nm-25). Normal and high light intensity at 100 and 500 μE was achieved by adjusting the distance between the growth chamber and the light source. ^13^CO_2_ labelling was started at dawn by supplying the growth chamber with a mixture of CO_2_ air and ^13^CO_2_ air at equal volume a continuous flow rate 6 L/min. The Arabidopsis plants were labelled for 2, 5 and 8 hours (T_2_, T_5_ and T_8_) before their shoots cut and snap-frozen in liquid nitrogen to stop all biological activities immediately. Three biological replicates were collected at each time point.

### Protein extraction, in-gel/solution digestion, high pH HPLC separation and LC-MS analysis of tryptic peptides

The shoot samples (~0.1 g) were snap-frozen in liquid nitrogen and homogenized using Qiagen tissue lysis beads (2 mm). A total plant protein extraction kit (PE0230-1KT, Sigma Chemicals) was used to extract total proteins. The final pellet of total protein was dissolved in Solution 4 and then reduced and alkylated by tributylphosphine (TBP) and iodoacetamide (IAA) as described in the Sigma manual. The suspension was centrifuged at 16,000 g for 30 min, and the supernatant was assayed for protein concentration by amido black quantification as described previously (Li et al., 2012). Protein (100 μg) in solution from each sample was then mixed with an equal volume of 2×sample buffer (4% SDS, 125 mM Tris, 20% glycerol, 0.005% Bromophenol blue and 10% mercaptoethanol, pH6.8) before being separated on a Biorad protean II electrophoresis system with a 4% (v/v) polyacrylamide stacking gel and 12% (v/v) polyacrylamide separation gel. Proteins were visualized by colloidal Coomassie Brilliant Blue G250 staining. One gel lane from one single biological replicate was excised into 11 fractions. Gel lanes from ten samples (T_0_ as a control, three biological replicates for ^13^C labelled sample at each time point) were fractioned into 110 and in-gel digested as described previously (Li et al., 2017). For protein abundance measurements, total proteins (50 μg) from ^14^N grown plant were combined with the fully ^15^N labelled protein reference (50 μg), and then in-solution trypsin digested. Each sample was separated into 96 fractions by high pH HPLC separation and further pooled into 6 fractions. Twenty-one total protein samples (Three biological samples from T_0_; 2, 5 and 8 under both standard and high light conditions kept in the same growth chamber as ^13^C labelling) were in-solution digested and separated into 126 fractions. Tryptic peptides from in-gel/in-solution digested were lyophilized in a Labconco centrifugal vacuum concentrator. Lyophilized samples were first resuspended in loading buffer (5% ACN, 0.% FA) and filtered through 0.22 μm Millipore column before being run in an Orbitrap Fusion (Thermo Fisher Scientific) mass spectrometer over the course of 95 mins over 2–30% (v/v) acetonitrile in 0.1% (v/v) formic acid (Dionex UltiMate 3000) on a 250 × 0.075 mm column (Dr. Maisch Reprosil-PUR 1.9 mm).

### Mass spectrometry data analysis

Orbitrap fusion raw (.raw) files were first converted to mzML using the Msconvert package from the Proteowizard project, and mzML files were subsequently converted to Mascot generic files (.mgf) using the mzxml2 search tool from the TPP. Mascot generic file peak lists were searched against an in-house *Arabidopsis* database comprising ATH1.pep (release 10) from The *Arabidopsis* Information Resource (TAIR) and the *Arabidopsis* mitochondrial and plastid protein sets (33621 sequences; 13487170 residues) (Lamesch et al., 2012), using the Mascot search engine version 2.3 and utilizing error tolerances of 10 ppm for MS and 0.5 Da for MS/MS; “Max Missed Cleavages” set to 1; variable modifications of oxidation (Met) and carbamidomethyl (Cys). All mzML files and dat files are provided in ProteomeXchange. We used iProphet and ProteinProphet from the Trans Proteomic Pipeline (TPP) to analyze peptide and protein probability and global false discovery rate (FDR) (Nesvizhskii et al., 2003; Deutsch et al., 2010; Shteynberg et al., 2011). The reported peptide lists with p=0.8 have FDRs of <3%, and protein lists with p=0.95 have FDRs of <0.5%. Quantification of LPFs (^13^C labeled protein fraction) and protein abundance (^14^N/^15^N ratios) were accomplished by an in-house script written in R as described previously (Nelson et al., 2014b; Li et al., 2017; Salih et al., 2020). Mass spectrometry data can be accessed through Proteomexchange through two entries: protein abundance changes in response to high light treatment (PXD010888), Protein turnover rates under high light treatment (PXD010889).

### Total RNA-sequencing

Arabidopsis plants were grown under identical conditions as per the ^13^C labelling experiment, except that normal air was supplied to the growth chamber. Shoot tissues were harvested (as above) in biological triplicate after differing light transitions from dark: T0D - end of night (dark control), T2H - 2 hours high light, T2L - 2 hours standard light, T5H - 5 hours high light, T5L - 5 hours standard light, T8H - 8 hours high light, and T8L - 8 hours standard light. Total RNA was isolated using TRI reagent based on an adapted protocol (Crisp et al., 2017). Full details are available at protocols.io: dx.doi.org/10.17504/protocols.io.bt8wnrxe. Total RNA-sequencing libraries were prepared using the TruSeq Stranded Total RNA with Ribo-Zero Plant kit (RS-122-2402, Illumina, CA, USA) as per manufacturer’s instructions but with input RNA and reaction volumes adjusted by one-third. PCR amplified libraries were pooled equal-molar and sequenced (75 bp, single-end) on one lane of the NextSeq500.

Raw read quality was first diagnosed using FastQC (v0.11.7). Trim Galore! (v0.4.4) was used for adapter and low-quality read trimming with PHRED score < 20 (-q 20). Trimmed reads were input for single-end splice-aware alignments using Subjunc from the Subread package v1.5.0 (Liao et al., 2013), retaining only reads that uniquely aligned to the Arabidopsis TAIR10 reference genome. Uniquely aligned reads were sorting and indexed using Samtools v1.3.1 (Li et al., 2009). Aligned reads were summarised to gene-level loci using the Araport11 annotation (Cheng et al., 2017) using featureCounts (-s 2 for reverse stranded libraries) (Liao et al., 2014)). Differential gene expression was tested using the edgeR quasi-likelihood pipeline(Robinson and Oshlack, 2010; Chen et al., 2016). Reads mapping to ribosomal RNA were removed; only loci containing counts per million (CPM) > 1 in at least three samples were examined. The trimmed mean of M-values (TMM) method was used for library normalisation to account for sequencing depth and composition (Robinson and Oshlack, 2010). Generalized linear models were fitted using normalized counts to estimate dispersion (glmQLFit) followed by employing quasi-likelihood F-tests (glmQLFTest) to test for differential expression while controlling for false discovery rates due to multiple hypothesis testing (FDR adjusted p-value < 0.05). Protein subcellular localisation data was acquired from SUBA (Hooper et al., 2017).

RNA-seq data is summarized in **DataS1** and can be accessed at GEO repository GSE131545: https://www.ncbi.nlm.nih.gov/geo/query/acc.cgi?acc=GSE131545. Code used for analyses are available on GitHub: https://github.com/dtrain16/NGS-scripts.

### PSII fluorescence parameters measured by mini-PAM and IMAGING-PAM

The Arabidopsis plants grown under the same condition for labelling experiment were used for PSII fluorescence parameters measurements except that normal air was supplied to the growth chamber. After darkness adaption at least 20 mins, PSII parameters of T_0_ plants were measured using LED as the light source at 100 and 500 μE light intensity with a mini-PAM (Heinz Walz GmbH). Fo/F values were measured every 1min for 1hour. Y(II), Y(NPQ), and Y(NO) values were calculated with the WinControl-3.25 data acquisition software. For Fv/Fm measurements, T_0_ and T_1_, T_2_, T_5_, and T_8_ plants light exposed at 100 and 500 μE were darkness adapted at least 20 mins before being measured by a MAXI version of the IMAGING-PAM (Heinz Walz GmbH). A color gradient was used to demonstrate the Fv/Fm (maximum quantum yield of PSII) values measured by IMAGING-PAM in leaves of the whole rosette. One biological replicate was a combination of measured Fv/Fm values in three leaves in Arabidopsis plants.

### Metabolite Extraction

The Arabidopsis plants grown under the same condition for labelling experiment were used for metabolite extraction except that normal air was supplied to the growth chamber. Plant tissues (15–50 mg) were collected at specified time points and immediately snap-frozen in liquid nitrogen. Samples were ground tofine powder and 500 μl of cold metabolite extraction solution (90% [v/v] methanol, spiked with 2 mg/ml ribitol, 6 mg/ml adipic acid, and 2 mg/ml and ^13^C-leucine as internal standards). Samples were immediately vortexed and shaken at 1,400 rpm for 20 min at 75°C. Cell debris was removed by centrifugation at 20,000 × g for 5 minutes. For each sample, 100 or 400 μl of supernatant was transferred to a new tube and either proceeded to derivatization for LC-MS analysis or dried using a SpeedVac.

### Analyses of organic acids and amino acids by selective reaction monitoring using triple quadrupole (QQQ) mass spectrometry

For LC-MS analysis of organic acids, sample derivatization was carried out based on previously published methods with modifications (Han et al., 2013). Briefly, for each of 100 μL of sample, 50 μL of 250 mM 3-nitrophenylhydrazine in 50% methanol, 50 μL of 150 mM 1-ethyl-3-(3-dimethylaminopropyl) carbodiimide in methanol, and 50 μL of 7.5% pyridine in 75% methanol were mixed and allowed to react on ice for 60 minutes. To terminate the reaction, 50 μL of 2 mg/mL butylated-hydroxytoluene in methanol was added, followed by the addition of 700 μL of water. Derivatized organic acids were separated on a Phenomenex Kinetex XB-C18 column (50 × 2.1mm, 5μm particle size) using 0.1% formic acid in water (solvent A) and methanol with 0.1% formic acid (solvent B) as the mobile phase. The elution gradient was 18% B at 1 min, 90% B at 10 min, 100% B at 11 min, 100% B at 12 min, 18% B at 13 min and 18% B at 20 min. The column flow rate was 0.3 mL/min and the column temperature was maintained at 40 °C. The QQQ-MS was operated in the negative ion mode with multiple reaction monitoring (MRM) mode.

## Supporting information

DataS5

DataS3

DataS2

DataS1

## Acknowledgements

Jacob Petereit and Sandra Kebler are thanked for advice on plant growth conditions and lighting the chamber with LEDs. We acknowledge the Biomolecular Resource Facility at the ANU for assistance with Illumina sequencing and computational resources provided by the National Computational Infrastructure, which is supported by the Australian Government. Mass spectrometry was performed on instruments managed by the WA Proteomics Facility as a node of Proteomics Australia and was supported by infrastructure funding from the Western Australian State Government in partnership with Bioplatforms Australia under the Commonwealth Government National Collaborative Research Infrastructure Strategy

## AUTHOR CONTRIBUTION

LL, BP, AHM designed the research; LL, KS and AWY performed plant growth and plant physiological parameter measurements; DRG and PAC performed RNA-sequencing. Mass spectrometry proteomics and analysis was performed by LL and OD. CPL performed metabolites extraction and mass spectrometry analysis. LL and AHM wrote the manuscript. All authors contribute to the writing and revision of the article.

## FUNDING

This work was supported through funding by the Australian Research Council (CE140100008, FL200100057) to AHM and BP, National Natural Science Foundation of China (31970294) and Tianjin NSF (19JCYBJC24100) to LL, CSIRO Synthetic Biology Future Science Platform to DRG.

## COMPETING INTERESTS

The authors declare that there are no competing interests associated with the manuscript.

## Supplemental Figures

**Fig S1.**
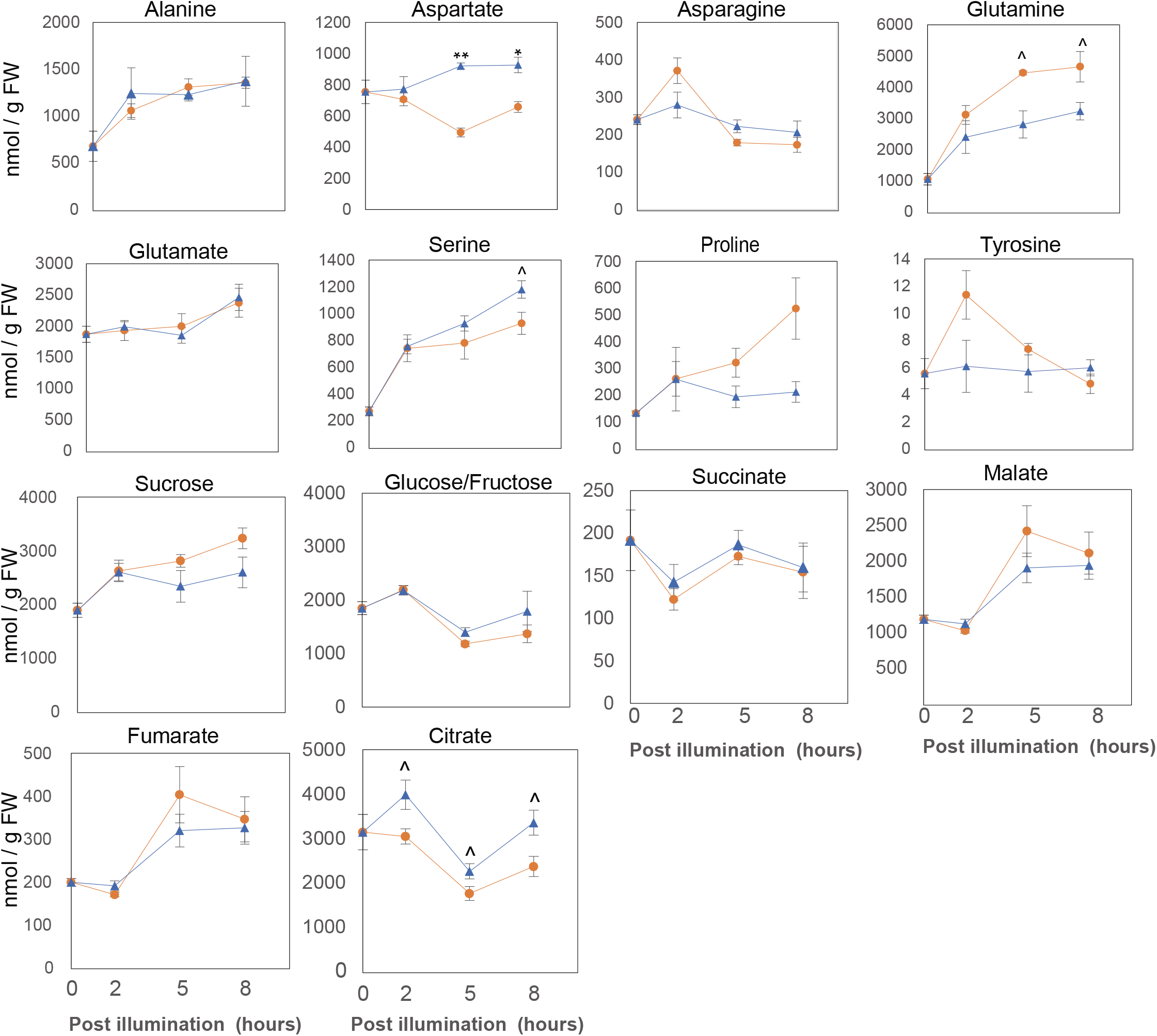
High light induced changes in abundance of specific amino acids and organic acids. Specific amino acids (measured by LC-QQQ MS) and organic acids (measured by LC-Q-TOF MS) that increase in abundance in response to high light treatment. Error bars show standard errors (biological replicates n=3). Statistical significance tests were performed with a student’s t test (**P<0.01, * P<0.05, ^ P<0.1).

**Fig S2.**
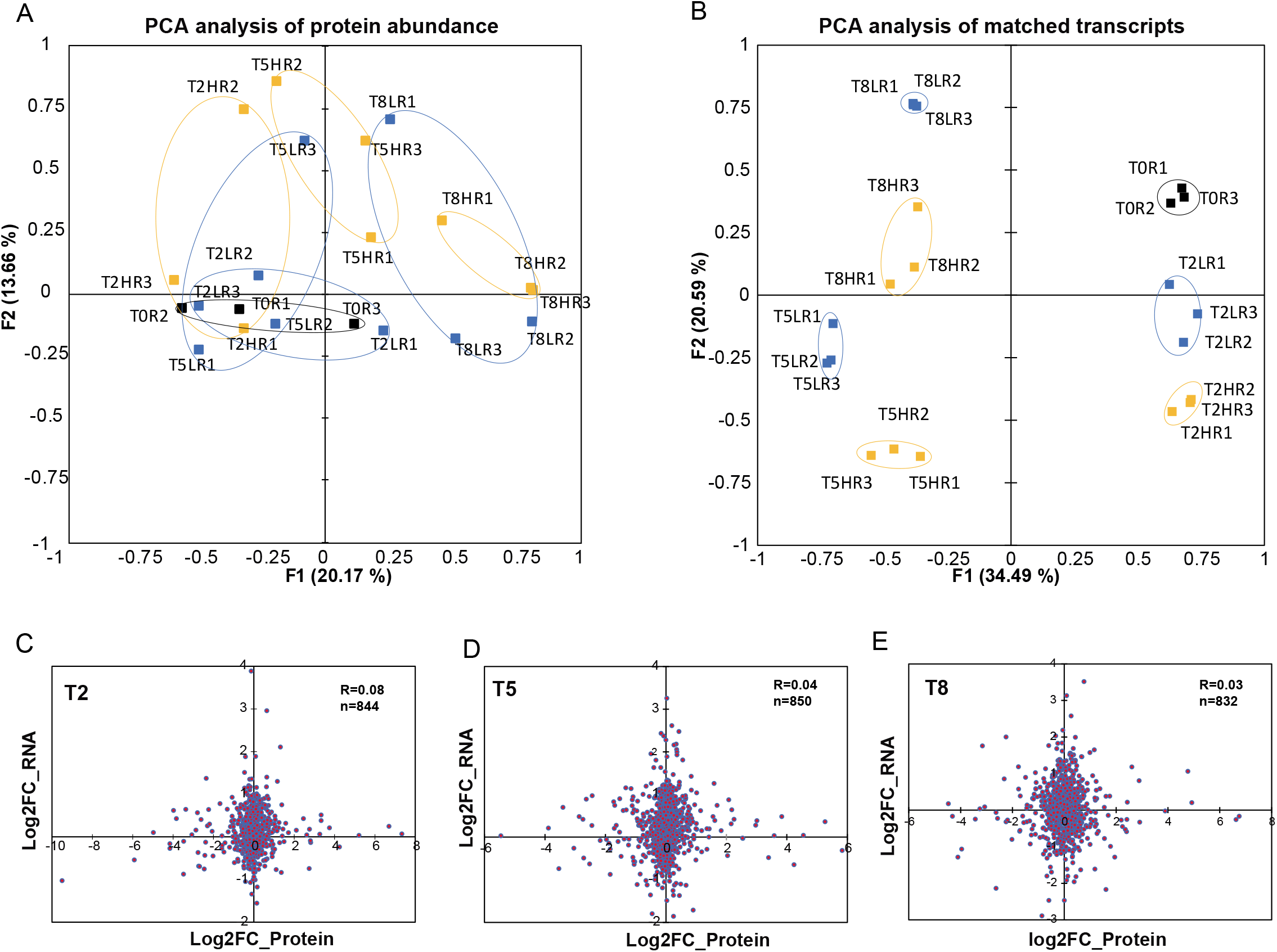
Principal components analysis and correlations of protein and mRNA abundance showing the relationship between transcriptional responses and protein abundance in standard light and high light conditions. PCA analysis for 370 proteins with measured transcript abundance (**DataS1**) and protein abundance (**DataS2**) in the dark (black), standard (blue), and high light (yellow) conditions. Scatterplots display the relationship between log2 fold-change in protein abundance (x-axis) and mRNA abundance (y-axis), in response to high light. Pearson’s *r* was calculated to quantify their correlation at 2h (T2), 5h (T5) and 8h (T8) (**C-E**).

**Fig S3.**
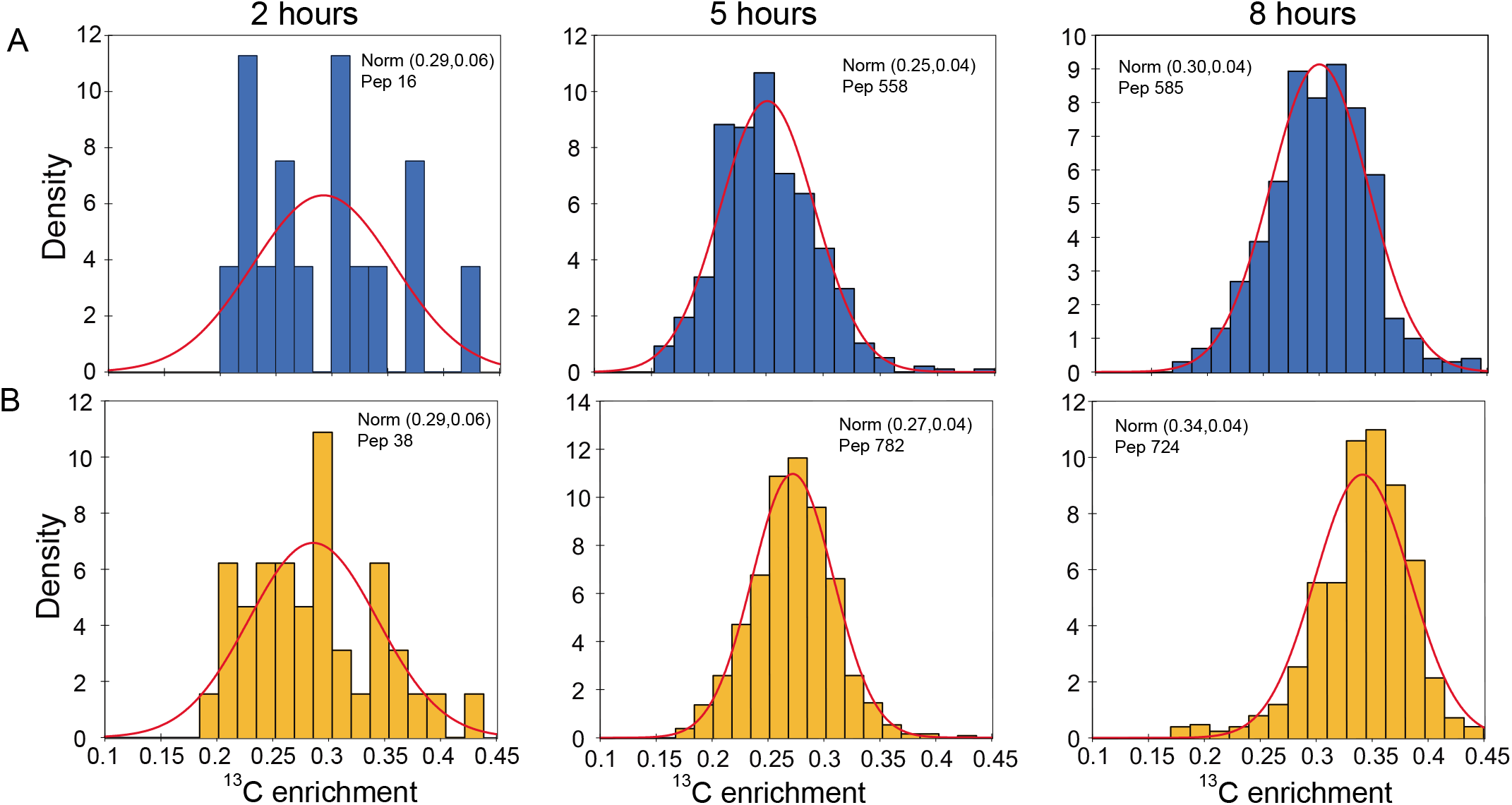
^13^C enrichment level of the heavy labelled ^13^C peptide population under standard (blue) and high light (yellow) conditions. The calculated ^13^C enrichment level for all peptides identified under each condition from the progressive labelling experiments combined. In each histogram, the bars are the actual peptide number, the median and standard deviation are shown as a plotted red line normal distribution (norm). The number of unique peptides (pep) included in each analysis is shown.

## Supplemental Data

**DataS1:** High light transcriptome analysis.

**DataS2:** High light proteome analysis.

**DataS3:** Protein turnover rates by ^13^C labelling in high and standard light.

**DataS4:** Comparison of protein turnover rates in standard and high light conditions.

**DataS5:** Changes in transcript and protein abundance of proteins with differences in protein turnover rate.

## Notes

### Competing Interest Statement

The authors have declared no competing interest.

## Reference

Aigner, H., Wilson, R.H., Bracher, A., Calisse, L., Bhat, J.Y., Hartl, F.U., and Hayer-Hartl, M. (2017). Plant RuBisCo assembly in E. coli with five chloroplast chaperones including BSD2. Science 358, 1272–1278.

Araújo, W.L., Tohge, T., Ishizaki, K., Leaver, C.J., and Fernie, A.R. (2011). Protein degradation – an alternative respiratory substrate for stressed plants. Trends Plant Sci.

Bhuiyan, N.H., Friso, G., Rowland, E., Majsec, K., and van Wijk, K.J. (2016). The Plastoglobule-Localized Metallopeptidase PGM48 Is a Positive Regulator of Senescence in Arabidopsis thaliana. Plant Cell 28, 3020–3037.

Boyes, D.C., Zayed, A.M., Ascenzi, R., McCaskill, A.J., Hoffman, N.E., Davis, K.R., and Gorlach, J. (2001). Growth stage-based phenotypic analysis of Arabidopsis: a model for high throughput functional genomics in plants. Plant Cell 13, 1499–1510.

Chatterjee, A., Abeydeera, N.D., Bale, S., Pai, P.J., Dorrestein, P.C., Russell, D.H., Ealick, S.E., and Begley, T.P. (2011). Saccharomyces cerevisiae THI4p is a suicide thiamine thiazole synthase. Nature 478, 542–546.

Chen, J.-H., Chen, S.-T., He, N.-Y., Wang, Q.-L., Zhao, Y., Gao, W., and Guo, F.-Q. (2020). Nuclear-encoded synthesis of the D1 subunit of photosystem II increases photosynthetic efficiency and crop yield. Nature Plants 6, 570–580.

Chen, Y., Lun, A.T., and Smyth, G.K. (2016). From reads to genes to pathways: differential expression analysis of RNA-Seq experiments using Rsubread and the edgeR quasi-likelihood pipeline. F1000Res 5, 1438.

Cheng, C.Y., Krishnakumar, V., Chan, A.P., Thibaud-Nissen, F., Schobel, S., and Town, C.D. (2017). Araport11: a complete reannotation of the Arabidopsis thaliana reference genome. Plant J 89, 789–804.

Chotewutmontri, P., and Barkan, A. (2016). Dynamics of Chloroplast Translation during Chloroplast Differentiation in Maize. PLoS Genet 12, e1006106.

Chotewutmontri, P., and Barkan, A. (2020). Light-induced psbA translation in plants is triggered by photosystem II damage via an assembly-linked autoregulatory circuit. Proceedings of the National Academy of Sciences 117, 21775–21784.

Collin, V., Issakidis-Bourguet, E., Marchand, C., Hirasawa, M., Lancelin, J.M., Knaff, D.B., and Miginiac-Maslow, M. (2003). The Arabidopsis plastidial thioredoxins: new functions and new insights into specificity. The Journal of biological chemistry 278, 23747–23752.

Crisp, P.A., Ganguly, D.R., Smith, A.B., Murray, K.D., Estavillo, G.M., Searle, I., Ford, E., Bogdanovic, O., Lister, R., Borevitz, J.O., Eichten, S.R., and Pogson, B.J. (2017). Rapid Recovery Gene Downregulation during Excess-Light Stress and Recovery in Arabidopsis. Plant Cell 29, 1836–1863.

Dai, S., Schwendtmayer, C., Schurmann, P., Ramaswamy, S., and Eklund, H. (2000). Redox signaling in chloroplasts: cleavage of disulfides by an iron-sulfur cluster. Science 287, 655–658.

Deutsch, E.W., Mendoza, L., Shteynberg, D., Farrah, T., Lam, H., Tasman, N., Sun, Z., Nilsson, E., Pratt, B., Prazen, B., Eng, J.K., Martin, D.B., Nesvizhskii, A.I., and Aebersold, R. (2010). A guided tour of the Trans-Proteomic Pipeline. Proteomics 10, 1150–1159.

Gabruk, M., and Mysliwa-Kurdziel, B. (2015). Light-Dependent Protochlorophyllide Oxidoreductase: Phylogeny, Regulation, and Catalytic Properties. Biochemistry 54, 5255–5262.

Gonzalez-Jorge, S., Ha, S.-H., Magallanes-Lundback, M., Gilliland, L.U., Zhou, A., Lipka, A.E., Nguyen, Y.-N., Angelovici, R., Lin, H., Cepela, J., Little, H., Buell, C.R., Gore, M.A., and DellaPenna, D. (2013). CAROTENOID CLEAVAGE DIOXYGENASE4 Is a Negative Regulator of β-Carotene Content in Arabidopsis Seeds. The Plant Cell 25, 4812–4826.

Guinea Diaz, M., Nikkanen, L., Himanen, K., Toivola, J., and Rintamäki, E. (2020). Two chloroplast thioredoxin systems differentially modulate photosynthesis in Arabidopsis depending on light intensity and leaf age. The Plant Journal 104, 718–734.

Guo, H., and Ecker, J.R. (2003). Plant responses to ethylene gas are mediated by SCF(EBF1/EBF2)-dependent proteolysis of EIN3 transcription factor. Cell 115, 667–677.

Gururani, M.A., Venkatesh, J., and Tran, L.S. (2015). Regulation of Photosynthesis during Abiotic Stress-Induced Photoinhibition. Mol Plant 8, 1304–1320.

Han, J., Gagnon, S., Eckle, T., and Borchers, C.H. (2013). Metabolomic analysis of key central carbon metabolism carboxylic acids as their 3-nitrophenylhydrazones by UPLC/ESI-MS. Electrophoresis 34, 2891–2900.

Hanson, A.D., McCarty, D.R., Henry, C.S., Xian, X., Joshi, J., Patterson, J.A., Garcia-Garcia, J.D., Fleischmann, S.D., Tivendale, N.D., and Millar, A.H. (2021). The number of catalytic cycles in an enzyme’s lifetime and why it matters to metabolic engineering. Proc Natl Acad Sci U S A 118.

Hildebrandt, T.M., Nunes Nesi, A., Araujo, W.L., and Braun, H.P. (2015). Amino Acid Catabolism in Plants. Mol Plant 8, 1563–1579.

Hoecker, U., and Quail, P.H. (2001). The phytochrome A-specific signaling intermediate SPA1 interacts directly with COP1, a constitutive repressor of light signaling in Arabidopsis. The Journal of biological chemistry 276, 38173–38178.

Holtorf, H., Reinbothe, S., Reinbothe, C., Bereza, B., and Apel, K. (1995). Two routes of chlorophyllide synthesis that are differentially regulated by light in barley (Hordeum vulgare L.). Proc Natl Acad Sci U S A 92, 3254–3258.

Hooper, C.M., Castleden, I.R., Tanz, S.K., Aryamanesh, N., and Millar, A.H. (2017). SUBA4: the interactive data analysis centre for Arabidopsis subcellular protein locations. Nucleic Acids Res 45, D1064–D1074.

Huang, J., Zhao, X., and Chory, J. (2019). The Arabidopsis Transcriptome Responds Specifically and Dynamically to High Light Stress. Cell Reports 29, 4186–4199.e4183.

Ishihara, H., Obata, T., Sulpice, R., Fernie, A.R., and Stitt, M. (2015). Quantifying Protein Synthesis and Degradation in Arabidopsis by Dynamic 13CO2 Labeling and Analysis of Enrichment in Individual Amino Acids in Their Free Pools and in Protein. Plant Physiol 168, 74–93.

Ishihara, H., Moraes, T.A., Pyl, E.T., Schulze, W.X., Obata, T., Scheffel, A., Fernie, A.R., Sulpice, R., and Stitt, M. (2017). Growth rate correlates negatively with protein turnover in Arabidopsis accessions. Plant J 91, 416–429.

Izumi, M., Ishida, H., Nakamura, S., and Hidema, J. (2017). Entire Photodamaged Chloroplasts Are Transported to the Central Vacuole by Autophagy. Plant Cell 29, 377–394.

Joshi, J., Beaudoin, G.A.W., Patterson, J.A., Garcia-Garcia, J.D., Belisle, C.E., Chang, L.Y., Li, L., Duncan, O., Millar, A.H., and Hanson, A.D. (2020). Bioinformatic and experimental evidence for suicidal and catalytic plant THI4s. Biochem J 477, 2055–2069.

Kato, Y., Ozawa, S., Takahashi, Y., and Sakamoto, W. (2015). D1 fragmentation in photosystem II repair caused by photo-damage of a two-step model. Photosynth Res 126, 409–416.

Kolling, K., George, G.M., Kunzli, R., Flutsch, P., and Zeeman, S.C. (2015). A whole-plant chamber system for parallel gas exchange measurements of Arabidopsis and other herbaceous species. Plant Methods 11, 48.

Kuhn, A., Engqvist, M.K.M., Jansen, E.E.W., Weber, A.P.M., Jakobs, C., Maurino, V.G., and Rennenberg, H. (2013). D-2-hydroxyglutarate metabolism is linked to photorespiration in theshm1-1mutant. Plant Biology 15, 776–784.

Lamesch, P., Berardini, T.Z., Li, D., Swarbreck, D., Wilks, C., Sasidharan, R., Muller, R., Dreher, K., Alexander, D.L., Garcia-Hernandez, M., Karthikeyan, A.S., Lee, C.H., Nelson, W.D., Ploetz, L., Singh, S., Wensel, A., and Huala, E. (2012). The Arabidopsis Information Resource (TAIR): improved gene annotation and new tools. Nucleic acids research 40, D1202–1210.

Lebedev, N., and Timko, M.P. (1999). Protochlorophyllide oxidoreductase B-catalyzed protochlorophyllide photoreduction in vitro: insight into the mechanism of chlorophyll formation in light-adapted plants. Proc Natl Acad Sci U S A 96, 9954–9959.

Li, H., Handsaker, B., Wysoker, A., Fennell, T., Ruan, J., Homer, N., Marth, G., Abecasis, G., Durbin, R., and Genome Project Data Processing, S. (2009). The Sequence Alignment/Map format and SAMtools. Bioinformatics 25, 2078–2079.

Li, L., Aro, E.M., and Millar, A.H. (2018). Mechanisms of Photodamage and Protein Turnover in Photoinhibition. Trends in plant science 23, 667–676.

Li, L., Nelson, C.J., Solheim, C., Whelan, J., and Millar, A.H. (2012). Determining degradation and synthesis rates of arabidopsis proteins using the kinetics of progressive 15N labeling of two-dimensional gel-separated protein spots. Molecular & cellular proteomics : MCP 11, M111 010025.

Li, L., Nelson, C.J., Trosch, J., Castleden, I., Huang, S., and Millar, A.H. (2017). Protein Degradation Rate in Arabidopsis thaliana Leaf Growth and Development. Plant Cell 29, 207–228.

Liang, C., Cheng, S.F., Zhang, Y.J., Sun, Y.Z., Fernie, A.R., Kang, K., Panagiotou, G., Lo, C., and Lim, B.L. (2016). Transcriptomic, proteomic and metabolic changes in Arabidopsis thaliana leaves after the onset of illumination. Bmc Plant Biol 16.

Liao, Y., Smyth, G.K., and Shi, W. (2013). The Subread aligner: fast, accurate and scalable read mapping by seed-and-vote. Nucleic Acids Res 41, e108.

Liao, Y., Smyth, G.K., and Shi, W. (2014). featureCounts: an efficient general purpose program for assigning sequence reads to genomic features. Bioinformatics 30, 923–930.

Ling, Q., Broad, W., Trösch, R., Töpel, M., Demiral Sert, T., Lymperopoulos, P., Baldwin, A., and Jarvis, R.P. (2019). Ubiquitin-dependent chloroplast-associated protein degradation in plants. Science 363.

Long, S.P., Humphries, S., and Falkowski, P.G. (1994). Photoinhibition of Photosynthesis in Nature. Annu Rev Plant Phys 45, 633–662.

Marri, L., Zaffagnini, M., Collin, V., Issakidis-Bourguet, E., Lemaire, S.D., Pupillo, P., Sparla, F., Miginiac-Maslow, M., and Trost, P. (2009). Prompt and Easy Activation by Specific Thioredoxins of Calvin Cycle Enzymes of Arabidopsis thaliana Associated in the GAPDH/CP12/PRK Supramolecular Complex. Molecular Plant 2, 259–269.

Matsumoto, F., Obayashi, T., Sasaki-Sekimoto, Y., Ohta, H., Takamiya, K., and Masuda, T. (2004). Gene expression profiling of the tetrapyrrole metabolic pathway in Arabidopsis with a mini-array system. Plant Physiol 135, 2379–2391.

Michaeli, S., Galili, G., Genschik, P., Fernie, A.R., and Avin-Wittenberg, T. (2016). Autophagy in Plants - What’s New on the Menu? Trends Plant Sci 21, 134–144.

Michelet, L., Zaffagnini, M., Morisse, S., Sparla, F., Perez-Perez, M.E., Francia, F., Danon, A., Marchand, C.H., Fermani, S., Trost, P., and Lemaire, S.D. (2013). Redox regulation of the Calvin-Benson cycle: something old, something new. Front Plant Sci 4, 470.

Millar, A.H., Heazlewood, J.L., Giglione, C., Holdsworth, M.J., Bachmair, A., and Schulze, W.X. (2019). The Scope, Functions, and Dynamics of Posttranslational Protein Modifications. Annu Rev Plant Biol 70, 119–151.

Muhlbauer, S.K., and Eichacker, L.A. (1998). Light-dependent formation of the photosynthetic proton gradient regulates translation elongation in chloroplasts. The Journal of biological chemistry 273, 20935–20940.

Nakai, M. (2018). New Perspectives on Chloroplast Protein Import. Plant and Cell Physiology 59, 1111–1119.

Nelson, C.J., Li, L., and Millar, A.H. (2014a). Quantitative analysis of protein turnover in plants. Proteomics 14, 579–592.

Nelson, C.J., Alexova, R., Jacoby, R.P., and Millar, A.H. (2014b). Proteins with high turnover rate in barley leaves estimated by proteome analysis combined with in planta isotope labeling. Plant Physiol 166, 91–108.

Nesvizhskii, A.I., Keller, A., Kolker, E., and Aebersold, R. (2003). A statistical model for identifying proteins by tandem mass spectrometry. Anal Chem 75, 4646–4658.

Nikkanen, L., and Rintamaki, E. (2019). Chloroplast thioredoxin systems dynamically regulate photosynthesis in plants. Biochem J 476, 1159–1172.

Nishimura, K., Kato, Y., and Sakamoto, W. (2016). Chloroplast Proteases: Updates on Proteolysis within and across Suborganellar Compartments. Plant Physiol 171, 2280–2293.

Ohad, I., Kyle, D.J., and Arntzen, C.J. (1984). Membrane protein damage and repair: removal and replacement of inactivated 32-kilodalton polypeptides in chloroplast membranes. J Cell Biol 99, 481–485.

Ohad, I., Kyle, D.J., and Hirschberg, J. (1985). Light-dependent degradation of the Q(B)-protein in isolated pea thylakoids. EMBO J 4, 1655–1659.

Priya, R., Sneha, P., Rivera Madrid, R., Doss, C.G.P., Singh, P., and Siva, R. (2017). Molecular Modeling and Dynamic Simulation of Arabidopsis Thaliana Carotenoid Cleavage Dioxygenase Gene: A Comparison with Bixa orellana and Crocus Sativus. Journal of Cellular Biochemistry 118, 2712–2721.

Robinson, M.D., and Oshlack, A. (2010). A scaling normalization method for differential expression analysis of RNA-seq data. Genome Biol 11, R25.

Sakuraba, Y., Rahman, M.L., Cho, S.-H., Kim, Y.-S., Koh, H.-J., Yoo, S.-C., and Paek, N.-C. (2013). The ricefaded green leaflocus encodes protochlorophyllide oxidoreductase B and is essential for chlorophyll synthesis under high light conditions. The Plant Journal 74, 122–133.

Salih, K.J., Duncan, O., Li, L., O’Leary, B., Fenske, R., Trosch, J., and Millar, A.H. (2020). Impact of oxidative stress on the function, abundance, and turnover of the Arabidopsis 80S cytosolic ribosome. Plant J 103, 128–139.

Schuster, M., Gao, Y., Schottler, M.A., Bock, R., and Zoschke, R. (2020). Limited Responsiveness of Chloroplast Gene Expression during Acclimation to High Light in Tobacco. Plant Physiol 182, 424–435.

Selinski, J., and Scheibe, R. (2019). Malate valves: old shuttles with new perspectives. Plant Biol (Stuttg) 21 Suppl 1, 21–30.

Shteynberg, D., Deutsch, E.W., Lam, H., Eng, J.K., Sun, Z., Tasman, N., Mendoza, L., Moritz, R.L., Aebersold, R., and Nesvizhskii, A.I. (2011). iProphet: multi-level integrative analysis of shotgun proteomic data improves peptide and protein identification rates and error estimates. Molecular & cellular proteomics : MCP 10, M111 007690.

Sundby, C., Mccaffery, S., and Anderson, J.M. (1993). Turnover of the Photosystem-Ii D1-Protein in Higher-Plants under Photoinhibitory and Nonphotoinhibitory Irradiance. Journal of Biological Chemistry 268, 25476–25482.

Tivendale, N.D., Fenske, R., Duncan, O., and Millar, A.H. (2021). In vivo homopropargylglycine incorporation enables sampling, isolation and characterization of nascent proteins from Arabidopsis thaliana. Plant J.

Trebitsh, T., and Danon, A. (2001). Translation of chloroplast psbA mRNA is regulated by signals initiated by both photosystems II and I. Proc Natl Acad Sci U S A 98, 12289–12294.

van Wijk, K.J. (2015). Protein maturation and proteolysis in plant plastids, mitochondria, and peroxisomes. Annu Rev Plant Biol 66, 75–111.

van Wijk, K.J., and Kessler, F. (2017). Plastoglobuli: Plastid Microcompartments with Integrated Functions in Metabolism, Plastid Developmental Transitions, and Environmental Adaptation. Annual Review of Plant Biology 68, 253–289.

Vitlin Gruber, A., and Feiz, L. (2018). Rubisco Assembly in the Chloroplast. Front Mol Biosci 5, 24.

Vogel, M.O., Moore, M., Konig, K., Pecher, P., Alsharafa, K., Lee, J., and Dietz, K.J. (2014). Fast retrograde signaling in response to high light involves metabolite export, MITOGEN-ACTIVATED PROTEIN KINASE6, and AP2/ERF transcription factors in Arabidopsis. Plant Cell 26, 1151–1165.

Wei, S.S., Niu, W.T., Zhai, X.T., Liang, W.Q., Xu, M., Fan, X., Lv, T.T., Xu, W.Y., Bai, J.T., Jia, N., and Li, B. (2019). Arabidopsis mtHSC70-1 plays important roles in the establishment of COX-dependent respiration and redox homeostasis. J Exp Bot 70, 5575–5590.

Woodson, J.D., Joens, M.S., Sinson, A.B., Gilkerson, J., Salome, P.A., Weigel, D., Fitzpatrick, J.A., and Chory, J. (2015). Ubiquitin facilitates a quality-control pathway that removes damaged chloroplasts. Science 350, 450–454.

Yu, J., Li, Y., Qin, Z., Guo, S., Li, Y., Miao, Y., Song, C., Chen, S., and Dai, S. (2020). Plant Chloroplast Stress Response: Insights from Thiol Redox Proteomics. Antioxid Redox Signal 33, 35–57.

Zaltsman, A., Feder, A., and Adam, Z. (2005). Developmental and light effects on the accumulation of FtsH protease in Arabidopsis chloroplasts--implications for thylakoid formation and photosystem II maintenance. Plant J 42, 609–617.

